# Dynamic pharmacophores unveil binding mode ensembles for classical partial agonists at the M_2_ receptor

**DOI:** 10.1101/2025.09.17.671681

**Authors:** Friederike Wunsch, Michael Kauk, Jenny Filor, Gerhard Wolber, Ulrike Holzgrabe, Carsten Hoffmann, Marcel Bermudez

**Affiliations:** Institute for Pharmaceutical and Medicinal Chemistry, University of Münster, Münster, Germany; Institute for Molecular Cell Biology, CMB-Center for Molecular Biomedicine, University Hospital Jena, Friedrich-Schiller University Jena, Jena, Germany; Department of Biology, Chemistry and Pharmacy, Institute for Pharmacy, Freie Universität Berlin, Berlin, Germany; Department of Pharmaceutical and Medicinal Chemistry, Institute of Pharmacy, University of Würzburg, Würzburg, Germany

**Author notes:** To whom correspondence should be addressed: Marcel Bermudez.

## Abstract

For the prototypic M_2_ receptor it has been previously demonstrated that dualsteric partial agonists can stabilize both, active and inactive receptor states, but it remains unclear whether orthosteric partial agonists have a similar mechanism. Here, we apply dynamic pharmacophores to unveil binding mode ensembles for classical M_2_ partial agonists. We report correlations between the spatial distribution of lipophilic contacts and ligand efficacy and demonstrate the applicability of dynamic pharmacophores to analyze subtle binding mode changes.

## Introduction

G protein-coupled receptors (GPCR) are integral membrane proteins that regulate a plethora of cellular functions. Their mechanistic complexity allows a fine-tuning of pharmacological responses beyond simple on-off-switches, which renders GPCRs a large and versatile drug target class.^1^ GPCR ligands can be classified based on their mechanistic effect and their resulting signaling response. This includes classical ligand categories like agonists, antagonists, or inverse agonists depending on their influence on basal receptor activity, but also allosteric modulators and biased ligands have been described.^2^ All GPCR ligands have in common that they interfere with the receptor’s conformational flexibility and thereby control binding of intracellular effector proteins. However, the underlying mechanisms are not yet fully understood. In particular, the degree of ligand-induced receptor activation can vary depending on the individual receptor-ligand pair.^2^ Full agonists result in 100% receptor activation, a value that is typically defined by endogenous ligands, e.g. acetylcholine for muscarinic receptors. While stronger receptor activation can be induced by a so-called superagonist, also graded receptor activation can be achieved by partial agonists. Classical examples for the prototypical muscarinic M_2_ receptor are iperoxo (IPX) as superagonist and pilocarpine as partial agonist.^3,4^

Different pharmacological, biophysical and computational approaches have been used to unveil the cause for different agonistic efficacies. Although two main concepts explain partial receptor activation, both suggested mechanisms might be applicable depending on the individual ligand-receptor pair. One major concept is the ligand-dependent stabilization of a distinct receptor conformation, often described as an intermediate conformation capable to induce an intracellular response that is weaker than for a full agonist.^5–8^ This phenomenon was also found to contain a kinetically driven component.^9–11^ Contrary to this explanation, there are several studies suggesting a shift in the equilibrium between active and inactive receptor states resulting in ligand-specific populations of functionally different receptor states.^12–16^

Previously, we have demonstrated that ligands can adopt different binding modes, which stabilize active, but also inactive receptor states of the M_2_ receptor.^17^ These dualsteric ligands were shown to bind to the allosteric vestibule and stabilize inactive receptor states in addition to their agonistic dualsteric binding mode. Ligand modifications and receptor mutagenesis indicated the possibility to fine-tune receptor activation from weak partial agonists (*isox-6-naph*) to full agonists (*iper-rigid-naph*). While this study showed the existence of binding mode ensembles for partial M_2_ receptor agonists, the special character of dualsteric ligands that are large and bind simultaneously at two binding sites remained a relevant limitation. We hypothesized that such binding mode ensembles also play a role for classical partial agonists that bind solely to the orthosteric binding site, but pinpointing subtle binding mode changes remained challenging.

To analyze and visualize binding mode ensembles for classical partial M_2_ receptor agonists like pilocarpine, we now apply dynamic pharmacophores (dynophores). This method is a fully automated combination of molecular dynamics (MD) simulations and three-dimensional pharmacophore models and provides information on both, the spatial and temporal distribution of receptor-ligand interactions (Figure 1).^18^ Dynophores have been recently applied to understand biased signaling^19,20^, subtype selectivity^21^, or mutational effects^22^. Here, we report binding mode ensembles of pilocarpine and a set of other partial M_2_ receptor agonists as the likely mechanism for their partial agonistic activity.

**Figure 1.**
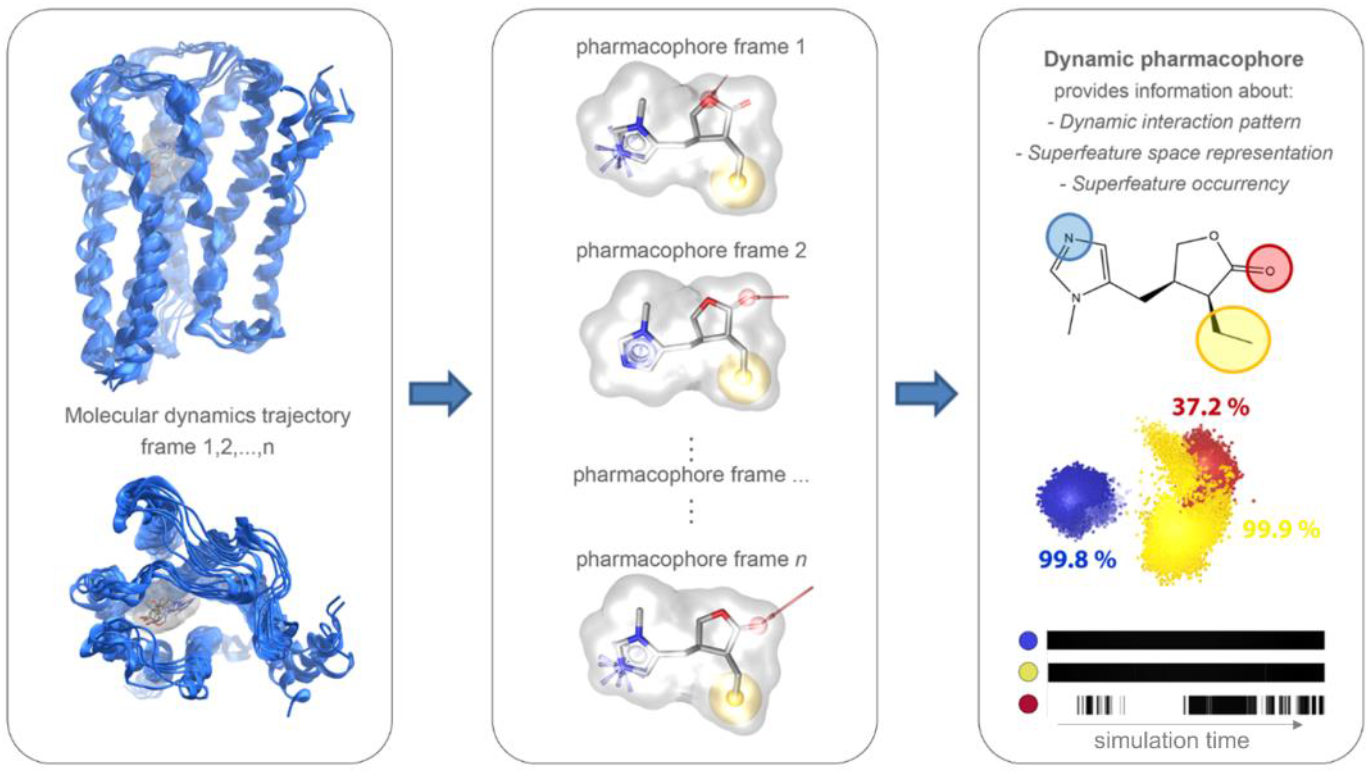
Workflow and output information for dynamic pharmacophores (dynophores). Based on classical all-atom MD simulation (left), static 3D-pharmacophores are generated for every single frame of the trajectory (middle), merged and grouped to superfeatures resulting in a dynamic pharmacophore model (right). Superfeatures are defined by feature type and involved atoms on the ligand side and account for the spatial distribution of receptor-ligand interactions. Barcodes indicate the temporal feature occurrence.

## Results and Discussion

Ligand-dependent receptor activation is characterized by a constriction of the transmembrane core region, in which the orthosteric binding pocket (OBP) is located (Figure S1).^17,23,24^ Binding site constriction is particularly pronounced for muscarinic receptors and was clearly demonstrated by crystal structures of the M_2_ receptor with bound inverse agonist quinuclidinylbenzilate (QNB) in comparison with the superagonist IPX.^1,23,25^ We run all-atoms MD simulations of those complexes, 1 µs each in triplicates and calculated dynophores. The ligand binding modes and respective receptor-ligand interactions showed only small variance as indicated by the sphere-like geometry of the interaction point clouds (Figure 2). The dynophore of IPX consists of only three spatially close features, underlining the stabilization of the active receptor conformation by a constricted OBP. In contrast, the two phenyl rings of QNB fill lipophilic subpockets, which interfere with the binding site constriction during receptor activation. A similar interaction pattern was found for the M_2_ inverse agonist *N*-methylscopolamine (NMS) that contains only one phenyl ring (Figure S2-S4). QNB and IPX show a highly stable binding mode and thereby effectively stabilize distinct conformations of the M_2_ receptor. However, other ligands including the partial agonist pilocarpine might show an ensemble of binding modes as previously reported for some dualsteric compounds^17^. To unveil such binding mode ensembles, we selected a set of partial agonists with varying agonist efficacies (Figure 3, Table S1): pilocarpine (E_max_ = 42%), alvameline (E_max_ = 49%), cevimeline (E_max_ = 84%), and talsaclidine (E_max_ = 92%). Agonist efficacies are not directly comparable, due to different assays used, but their rank order for agonist efficacy at M_2_ receptors is pilocarpine < alvameline < cevimeline < talsaclidine.

**Figure 2.**
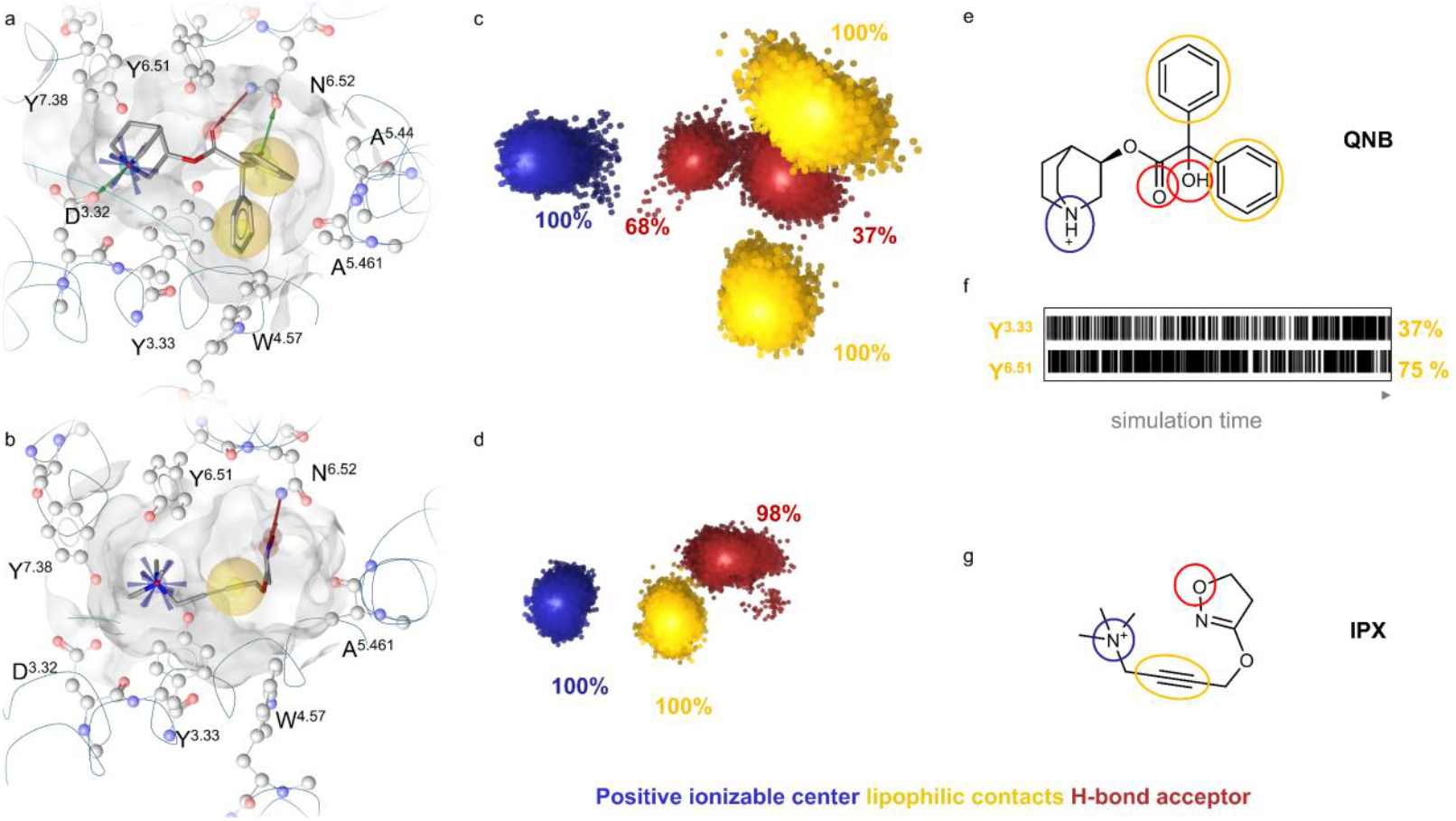
Dynophores indicate distinct interaction patterns for inverse agonists and full agonists. Static pharmacophores of (**a**) the inverse agonist QNB and (**b**) the superagonist iperoxo (IPX) bound to the M_2_ receptor. Space filling lipophilic groups of QNB stabilize inactive receptor conformations, while IPX allows for the contraction of the OBP finally stabilizing the active M_2_ receptor. Dynamic pharmacophores for QNB (**c, e**) and IPX (**d, g**) indicate stable, sphere-like interaction patterns. Percentages are mean values for feature occurrence in triplicate simulations (3x 1µs). Barcodes of important lipophilic contact residues underline these stable interactions for QNB (**f**)

**Figure 3.**
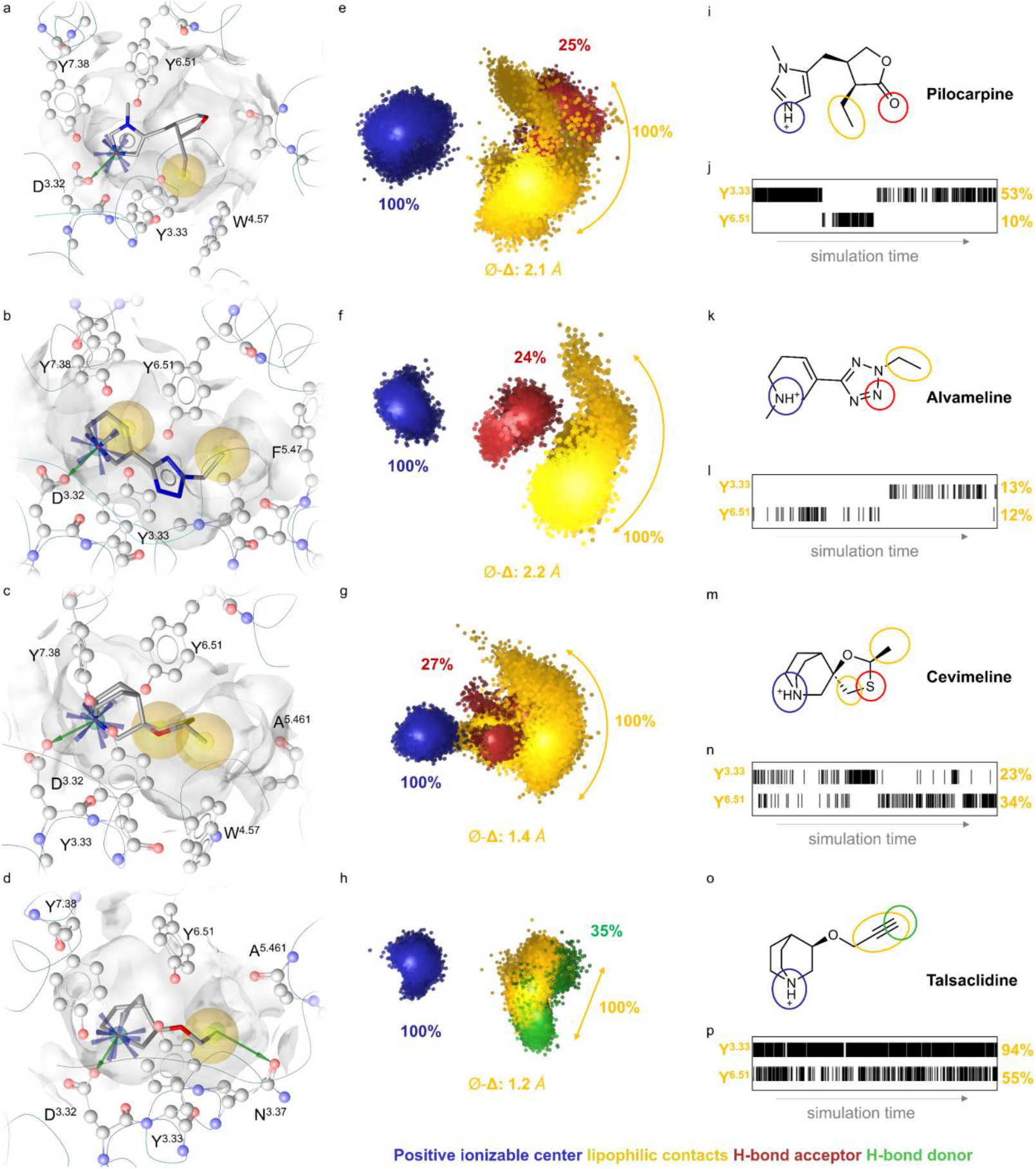
Dynamic interaction patterns of partial agonists correlate with reported agonist efficacy. Static pharmacophores of pilocarpine (**a**), alvameline (**b**), cevimeline (**c**), and talsaclidine (**d**) in the orthosteric binding pocket of the M_2_ receptor. The ethyl-γ-lactone of pilocarpine and the ethyl-tetrazole moiety of alvameline are sterically more demanding compared to the lipophilic substructures of cevimeline and talsaclidine. Dynophores of pilocarpine (**e**), alvameline (**f**), cevimeline (**g**) and talsaclidine (**h**). While the positive ionizable group of all partial agonists is fixed, the lipophilic substructure shows a half-moon shaped interaction pattern which gets narrower with increasing agonistic efficacy. ⌀-Δ defines the average distance of feature points to their centroid. Percentages are mean values for the feature occurrence in triplicate simulations (3x 1µs). 2D structures with highlighted interacting substructures (**I, k, m, o**). Barcodes of selected lipophilic contact residues for pilocarpine (**j**), alvameline (l), cevimeline (**n**) and talsaclidine (**p**) underline the switch between interactions with Y^3.33^ and Y^6.51^.

Like IPX and QNB, the dynophore analysis of all partial agonists indicated 100% occurrence of the positive ionizable feature in a condensed and sphere-like distribution. However, the small lipophilic moiety of each compound showed different orientations resulting in an ensemble of binding modes (Figure 3, Figure S3). Interestingly, the small lipophilic groups seem to occupy both lipophilic pockets also occupied by the phenyl moieties of QNB, but to different extents. Thereby, the dynamic spatial demand of each small lipophilic group correlates with the agonistic efficacy of the studied partial agonists (Figure S5, S6).

For the partial agonists with the lower agonistic efficacy (pilocarpine and alvameline), we observed two clearly distinguishable orientations of the ethyl group (Figure 3, Figure S7). For alvameline the scattering of the lipophilic feature was more variable and favoring the downward orientation compared to pilocarpine. Cevimeline and talsaclidine are less dynamic regarding the orientation of the small lipophilic moiety and therefore binding mode changes are visually not separable (Figure S8). Cevimeline, which has the most rigid scaffold, leads to an even distribution of the lipophilic feature with more narrowly distributed feature points compared to pilocarpine and alvameline. Talsaclidine, the partial agonist with the highest E_max_ among the studied ligands, indicates a narrow distribution of the lipophilic feature and does not point to the lipophilic pockets that hinder binding site constriction.

A comparison of the time-dependent interaction feature occurrence shows distinct changes between two binding modes for pilocarpine and alvameline, (Figure 3, Figure S7). Y^3.33^ and Y^6.51^ are located at opposite sites in the OBP and don’t interact simultaneously with the ligands. For pilocarpine, we see binding mode changes in both directions, which correlate with slightly staggered changes in distances measured throughout the OPB (Figure S7). Caused by a weaker interference with M_2_’s OBP contraction, the partial agonists cevimeline and talsaclidine show an increased simultaneity in the interactions with Y^3.33^ and Y^6.51^. The spatial proximity of the different binding modes of both ligands allows more fluid and subtle changes in the interacting residues (Figure 3, Figure S8).

To analyze the ligand’s impact on the binding site, we measured the sum of two distances as a surrogate parameter for OBP contraction (Figure S9). As expected, we observe larger distances for the inverse agonists (QNB, NMS) than for the super agonist IPX. Partial agonists indicate distance values in between. Unlike QNB, NMS and IPX the distance distribution for the partial agonists is much wider and not bell-shaped indicating a more heterogenous stabilization of OBP conformations.

Previous studies proved the concept of binding mode ensembles as mechanistic basis for partial receptor activation by using a series of dualsteric ligands, extensive pharmacological characterization and site-directed mutagenisis.^13,17^ In those studies, the dualsteric nature of the investigated ligands was essential to discriminate between functionally different binding modes, because different, overlapping binding sites were addressed. Although challenging, we now wanted to experimentally support the concept of binding mode ensembles for small organic partial agonists that bind to the receptor’s OBP. Therefore, we made use of alcuronium, an allosteric modulator that was found to preshape the receptor in an inactive state and should shift the binding mode ensemble of pilocarpine towards inactive receptor states. In line with our hypothesis, we observe a decrease in pilocarpine’s agonist activity when combined with alcuronium, but no change in agonist activity when combined with LY2119620 (Figure 4, Figure S10-11). The latter substance is a positive allosteric modulator that shows cooperativity with iperoxo.^23^

**Figure 4.**
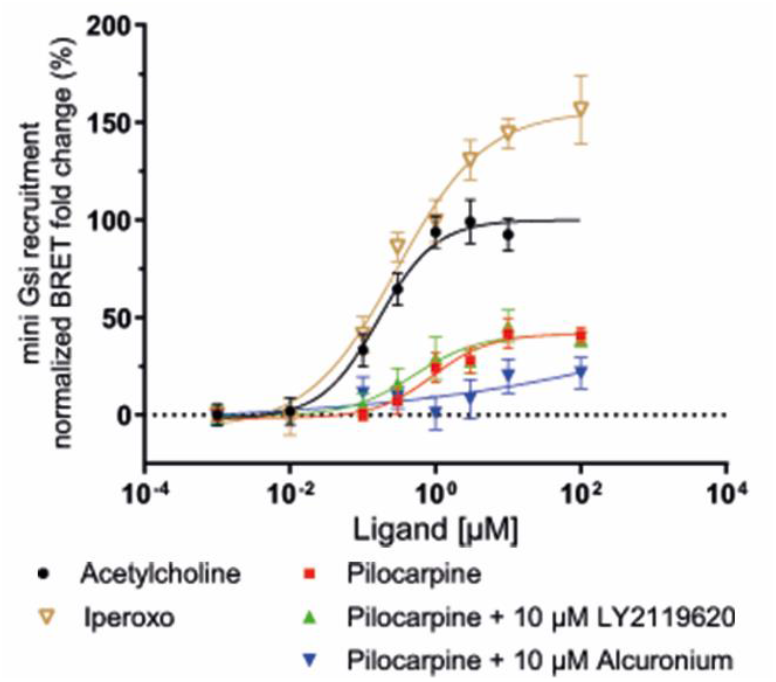
Concentration–response curves of the full agonist acetylcholine, the superagonist Iperoxo and the partial agonist pilocarpine at the M_2_ receptor using a BRET assay for miniG-protein recruitment. Combination of pilocarpine with the positive allosteric modulator LY2119620 showed no influence on pilocarpine’s agonist activity, but the negative allosteric modulator Alcuronium reduced pilocarpine’s agonist activity as expected. Each concentration was determined in triplicates, and each concentration– response curve was measured in three independent experiments.

While in general, binding mode ensembles might be highly abundant, there is only a small number of studies which link divergent binding modes to functionally different effects. For GPCRs, besides dualsteric muscarinic agonists^13,17^, a recent study was reported for cannabinoid receptors^15^. Other examples for which structural evidence was provided, including estrogen receptor α, farnesyltransferase, or 5-HT_3_ channels, are illustrated in Figure S12. We surmise that ligand binding mode ensembles are frequently occurring phenomena, that contribute to a specific pharmacological outcome in many cases. While X-ray crystallography, or cryo-EM might indicate multiple ligand binding modes, e.g. different ligand orientation within the binding site, it remains unclear whether smaller, but relevant differences can be structurally determined. Here, we have demonstrated the applicability of dynophore analysis to unveil subtle changes in binding modes of partial M_2_ receptor agonists. Moreover, we have linked distinct binding modes to specific interference with conformational changes during receptor activation and thereby support one possible mechanism for partial agonism at a prototypical GPCR.

## References

(1) Bock, A.; Bermudez, M. Allosteric coupling and biased agonism in G protein-coupled receptors. The FEBS journal 2021, 288 (8), 2513–2528. DOI: 10.1111/febs.15783.

(2) Hilger, D. The role of structural dynamics in GPCR-mediated signaling. The FEBS journal 2021, 288 (8), 2461–2489. DOI: 10.1111/febs.15841.

(3) Heinz, C. S.; Bermudez, M.; Jaiswal, N.; Große, C.; Kauk, M.; Hoffmann, C.; Holzgrabe, U. Hybridization into a Bitopic Ligand Increased Muscarinic Receptor Activation for Isopilocarpine but Not for Pilocarpine Derivatives. Journal of Natural Products 2023, 86 (4), 869–881. DOI: 10.1021/acs.jnatprod.2c01079.

(4) Schrage, R.; Seemann, W. K.; Klöckner, J.; Dallanoce, C.; Racké, K.; Kostenis, E.; Amici, M. de; Holzgrabe, U.; Mohr, K. Agonists with supraphysiological efficacy at the muscarinic M2 ACh receptor. British Journal of Pharmacology 2013, 169 (2), 357–370. DOI: 10.1111/bph.12003.

(5) Eddy, M. T.; Martin, B. T.; Wüthrich, K. A(2A) Adenosine Receptor Partial Agonism Related to Structural Rearrangements in an Activation Microswitch. Structure (London, England : 1993) 2021, 29 (2), 170-176.e3. DOI: 10.1016/j.str.2020.11.005.

(6) Masureel, M.; Zou, Y.; Picard, L.-P.; van der Westhuizen, E.; Mahoney, J. P.; Rodrigues João P.G.L.M.; Mildorf, T. J.; Dror, R. O.; Shaw, D. E.; Bouvier, M.; Pardon, E.; Steyaert, J.; Sunahara, R. K.; Weis, W. I.; Zhang, C.; Kobilka, B. K. Structural insights into binding specificity, efficacy and bias of a β2AR partial agonist. Nature chemical biology 2018, 14 (11), 1059–1066. DOI: 10.1038/s41589-018-0145-x.

(7) Zhang, Y.; Yang, F.; Ling, S.; Lv, P.; Zhou, Y.; Fang, W.; Sun, W.; Zhang, L.; Shi, P.; Tian, C. Single-particle cryo-EM structural studies of the β2AR–Gs complex bound with a full agonist formoterol. Cell Discovery 2020, 6 (1), 45. DOI: 10.1038/s41421-020-0176-9.

(8) Warne, T.; Edwards, P. C.; Doré, A. S.; Leslie, A. G. W.; Tate, C. G. Molecular basis for high-affinity agonist binding in GPCRs. Science (New York, N.Y.) 2019, 364 (6442), 775–778. DOI: 10.1126/science.aau5595.

(9) Nikolaev, V. O.; Hoffmann, C.; Bünemann, M.; Lohse, M. J.; Vilardaga, J.-P. Molecular basis of partial agonism at the neurotransmitter alpha2A-adrenergic receptor and Gi-protein heterotrimer. Journal of Biological Chemistry 2006, 281 (34), 24506–24511. DOI: 10.1074/jbc.M603266200.

(10) Gregorio, G. G.; Masureel, M.; Hilger, D.; Terry, D. S.; Juette, M.; Zhao, H.; Zhou, Z.; Perez-Aguilar, J. M.; Hauge, M.; Mathiasen, S.; Javitch, J. A.; Weinstein, H.; Kobilka, B. K.; Blanchard, S. C. Single-molecule analysis of ligand efficacy in β2AR–G-protein activation. Nature 2017, 547 (7661), 68–73. DOI: 10.1038/nature22354.

(11) Guo, D.; Mulder-Krieger, T.; IJzerman, A. P.; Heitman, L. H. Functional efficacy of adenosine A2A receptor agonists is positively correlated to their receptor residence time. British Journal of Pharmacology 2012, 166 (6), 1846–1859. DOI: 10.1111/j.1476-5381.2012.01897.x.

(12) Kofuku, Y.; Ueda, T.; Okude, J.; Shiraishi, Y.; Kondo, K.; Maeda, M.; Tsujishita, H.; Shimada, I. Efficacy of the β2-adrenergic receptor is determined by conformational equilibrium in the transmembrane region. Nature communications 2012, 3 (1), 1045. DOI: 10.1038/ncomms2046.

(13) Bock, A.; Chirinda, B.; Krebs, F.; Messerer, R.; Bätz, J.; Muth, M.; Dallanoce, C.; Klingenthal, D.; Tränkle, C.; Hoffmann, C.; Amici, M. de; Holzgrabe, U.; Kostenis, E.; Mohr, K. Dynamic ligand binding dictates partial agonism at a G protein-coupled receptor. Nature chemical biology 2014, 10 (1), 18– 20. DOI: 10.1038/nchembio.1384. Published Online: Nov. 10, 2013.

(14) Solt, A. S.; Bostock, M. J.; Shrestha, B.; Kumar, P.; Warne, T.; Tate, C. G.; Nietlispach, D. Insight into partial agonism by observing multiple equilibria for ligand-bound and Gs-mimetic nanobody-bound β1-adrenergic receptor. Nature communications 2017, 8 (1), 1795. DOI: 10.1038/s41467-017-02008-y.

(15) Dutta, S.; Selvam, B.; Das, A.; Shukla, D. Mechanistic origin of partial agonism of tetrahydrocannabinol for cannabinoid receptors. Journal of Biological Chemistry 2022, 298 (4), 101764. DOI: 10.1016/j.jbc.2022.101764.

(16) Claff, T.; Mahardhika, A. B.; Vaaßen, V. J.; Schlegel, J. G.; Vielmuth, C.; Weiße, R. H.; Sträter, N.; Müller, C. E. Structural Insights into Partial Activation of the Prototypic G Protein-Coupled Adenosine A2A Receptor. ACS Pharmacology & Translational Science 2024, 7 (5), 1415–1425. DOI: 10.1021/acsptsci.4c00051.

(17) Bock, A.; Bermudez, M.; Krebs, F.; Matera, C.; Chirinda, B.; Sydow, D.; Dallanoce, C.; Holzgrabe, U.; Amici, M. de; Lohse, M. J.; Wolber, G.; Mohr, K. Ligand Binding Ensembles Determine Graded Agonist Efficacies at a G Protein-coupled Receptor. Journal of Biological Chemistry 2016, 291 (31), 16375–16389. DOI: 10.1074/jbc.M116.735431.

(18) Schaller, D.; Šribar, D.; Noonan, T.; Deng, L.; Nguyen, T. N.; Pach, S.; Machalz, D.; Bermudez, M.; Wolber, G. Next generation 3D pharmacophore modeling. WIREs Comput Mol Sci 2020, 10 (4), e1468. DOI: 10.1002/wcms.1468.

(19) Denzinger, K.; Nguyen, T. N.; Noonan, T.; Wolber, G.; Bermudez, M. Biased Ligands Differentially Shape the Conformation of the Extracellular Loop Region in 5-HT(2B) Receptors. International journal of molecular sciences 2020, 21 (24). DOI: 10.3390/ijms21249728.

(20) Holze, J.; Bermudez, M.; Pfeil, E. M.; Kauk, M.; Bödefeld, T.; Irmen, M.; Matera, C.; Dallanoce, C.; Amici, M. de; Holzgrabe, U.; König, G. M.; Tränkle, C.; Wolber, G.; Schrage, R.; Mohr, K.; Hoffmann, C.; Kostenis, E.; Bock, A. Ligand-Specific Allosteric Coupling Controls G-Protein-Coupled Receptor Signaling. ACS Pharmacology & Translational Science 2020, 3 (5), 859–867. DOI: 10.1021/acsptsci.0c00069.

(21) Wunsch, F.; Nguyen, T. N.; Wolber, G.; Bermudez, M. Structural determinants of sphingosine-1-phosphate receptor selectivity. Archiv der Pharmazie 2023, 356 (12), e2300387. DOI: 10.1002/ardp.202300387.

(22) Wojciechowski, M. N.; Jokiel, J.; Kuss, H.; Bermúdez, M.; Jose, J. Combination of Autodisplay and Dynamic Pharmacophore Modeling Reveals New Insights into Cyclic Nucleotide Binding in Hyperpolarization-Activated and Cyclic Nucleotide-Gated Ion Channel 4 (HCN4). ACS Pharmacology & Translational Science 2024, 7 (12), 4010–4020. DOI: 10.1021/acsptsci.4c00497.

(23) Kruse, A. C.; Ring, A. M.; Manglik, A.; Hu, J.; Hu, K.; Eitel, K.; Hübner, H.; Pardon, E.; Valant, C.; Sexton, P. M.; Christopoulos, A.; Felder, C. C.; Gmeiner, P.; Steyaert, J.; Weis, W. I.; Garcia, K. C.; Wess, J.; Kobilka, B. K. Activation and allosteric modulation of a muscarinic acetylcholine receptor. Nature 2013, 504 (7478), 101–106. DOI: 10.1038/nature12735. Published Online: Nov. 20, 2013.

(24) Bermudez, M.; Bock, A.; Krebs, F.; Holzgrabe, U.; Mohr, K.; Lohse, M. J.; Wolber, G. Ligand-Specific Restriction of Extracellular Conformational Dynamics Constrains Signaling of the M2 Muscarinic Receptor. ACS Chemical Biology 2017, 12 (7), 1743–1748. DOI: 10.1021/acschembio.7b00275.

(25) Haga, K.; Kruse, A. C.; Asada, H.; Yurugi-Kobayashi, T.; Shiroishi, M.; Zhang, C.; Weis, W. I.; Okada, T.; Kobilka, B. K.; Haga, T.; Kobayashi, T. Structure of the human M2 muscarinic acetylcholine receptor bound to an antagonist. Nature 2012, 482 (7386), 547–551. DOI: 10.1038/nature10753. Published Online: Jan. 25, 2012.

