## Supplemental Information for "Dynamic pharmacophores unveil binding mode ensembles for classical partial agonists at the M_2_ receptor"

### Methods

#### Structure preparation and docking

We prepared all structural models in molecular operating environment v2020.0901 (MOE)[1] by adding missing sidechains and eliminating atom clashes. Ligands were built and protonated in MOE. PDB entry 4MQS (active $M_2$-iperoxo-Nb98)[2] was used for docking partial agonists in GOLD5.2[3] applying default settings and goldscore[4,5] as primary scoring function. The sidechain carbonyl of $D103^{3.32}$ with a radius of 15 Å was used to define the binding site. Binding poses were selected depending on static pharmacophore interactions as calculated in LigandScout4.4.3[6,7].

PDB entries 3UON (inactive $M_2$-QNB-T4L)[8] and 5ZKC (inactive $M_2$-NMS-BRIL)[9] were used for calculating static pharmacophores in inactive $M_2$. 5ZKC was used for docking Alcuronium. The oxygen atom of the phenolic hydroxyl group of $Y83^{2.63}$ with a radius of 15 Å was used to define the binding site. The sidechains of $E175^{ECL2}$, $Y177^{ECL2}$ and $W422^{7.34}$ were manually adapted.

#### Molecular dynamics simulations and analysis

While BRIL (thermostabilized apocytochrome b562 from *Escherichia coli* M7W/H102I/R106L) and T4L (T4 lysozym fusion protein) were removed from inactive $M_2$ structures, the nanobody Nb98 was maintained to stabilize the active $M_2$ conformation. Molecular dynamics simulations were prepared using Maestro v.12.7.156.[10] The complexes were titrated to pH7 and solvated in SPC water[11] containing 0.15 M NaCl. We placed the system in an orthorhombic box and applied periodic boundary conditions. $M_2$ was placed in a palmitoyl-oleoyl-phosphatidyl-choline (POPC) bilayer according to the orientations of proteins in membranes database (OPM)[12] entries of 3UON and 4MQS. The OPLS-AA force field[13] was used to parameterize the system.

Molecular dynamics simulations were calculated with Desmond v6.5.[14] on local GPUs (RTXA5000) for 1 µs in triplicates generating 20,000 frames as output. Throughout the simulations the number of particles, the temperature (300 K) and the pressure (atmospheric pressure) were kept constant.

Obtained trajectories were analyzed using VMD1.9.4[15]. The GPCRs backbones heavy atoms were aligned to those of frame 1. Atomic deviations are reported in tables S2 and S3. Dynamic pharmacophores were generated with the dynophore application[16] as integrated in the framework of LigandScout4.4.3.[7,17] For barcode representations as shown in figure 2, 3, S2, S4, S7, S8 every 40$^{th}$ frame of the trajectory was used. For the time-dependent color-coding of the lipophilic feature points and the distance measurements to the feature centroid, the feature coordinates were extracted from the dynophore file in LigandScout. The average distance between each feature point and the feature centroid was calculated in Excel. The file was converted to a CSV file. Using ScikitLearn[18], the points were displayed in a Cartesian coordinate system and color-coded according to their sequence in the CSV file.

**In vitro experiments**

The BRET assay for miniG-protein recruitment was essentially performed as previously described by Heinz et al.[19] and Leonhardt et al.[20]. In brief, HEK293T cells (originally obtained from DSMZ, Germany; ACC 305) were cultured at 37 °C, with 5% (v/v) $CO_2$ in Dulbecco's Modified Eagle Medium (DMEM, high glucose; Sigma-Aldrich, D6429), supplemented with 10% FBS, 0.1 mg/mL streptomycin, and 100 U/ml penicillin. The HEK293T cells were transiently transfected with FLAG-MR2-NanoLuc and the respective miniG$_{si}$-Venus, in a ratio of BRET donor and acceptor 1:2, using polyethylenimine (PEI MAX) reagent (Polysciences, 24765). PEI MAX was 2.5 times the amount of total DNA and PEI MAX–DNA mixture was solved in Opti-MEM (reduced serum medium, Gibco, 31985-070). After 24 h incubation at 37 °C, 5% (v/v) $CO_2$, cells were seeded on Poly-D-Lysine coated 96-well plates (BRAND, #781965) with 40,000 cells per well. The following day, immediately before measurement, cells were washed twice with measuring buffer (140 mM NaCl, 10 mM HEPES, 5.4 mM KCl, 2 mM $CaCl_2$, 1 mM $MgCl_2$; pH 7.3), followed by preincubation with 90 µL of measuring buffer supplemented with NanoLuc substrate (final dilution 1:3,500, Promega, N157B) at 37 °C for 1 min.

Measurements were performed with a Synergy Neo2 plate reader (BioTek/Agilent, Gen5 software) equipped with a Dual BRET2 PMT filter cube (400/510, BioTek, 1035072). After recording basal readings for five cycles (43 s interval, with an integration time of 0.3 s), 10 µL of measuring buffer supplemented with vehicle or different concentration of ligands was added manually. After ligand addition, the BRET signal was recorded with the same settings as for basal readings for 10 more cycles per plate. The BRET ratio (Venus/NanoLuc emission) was calculated by dividing averaged stimulated values over averaged basal readings and was normalized to vehicle stimulation. Each concentration was determined in triplicates, and each concentration–response curve was measured in at least three independent experiments.

The BRET data were analyzed by computer-aided nonlinear regression analysis using GraphPad Prism 10.4.2 software. The data set of each experiment was fitted by a four-parameter model with the Hill slope not higher than 1.1. Graphs were normalized to $E_{max}$ value for acetylcholine (100%), as reference ligand. The normalized data were plotted and fitted in GraphPad Prism 10 as described.

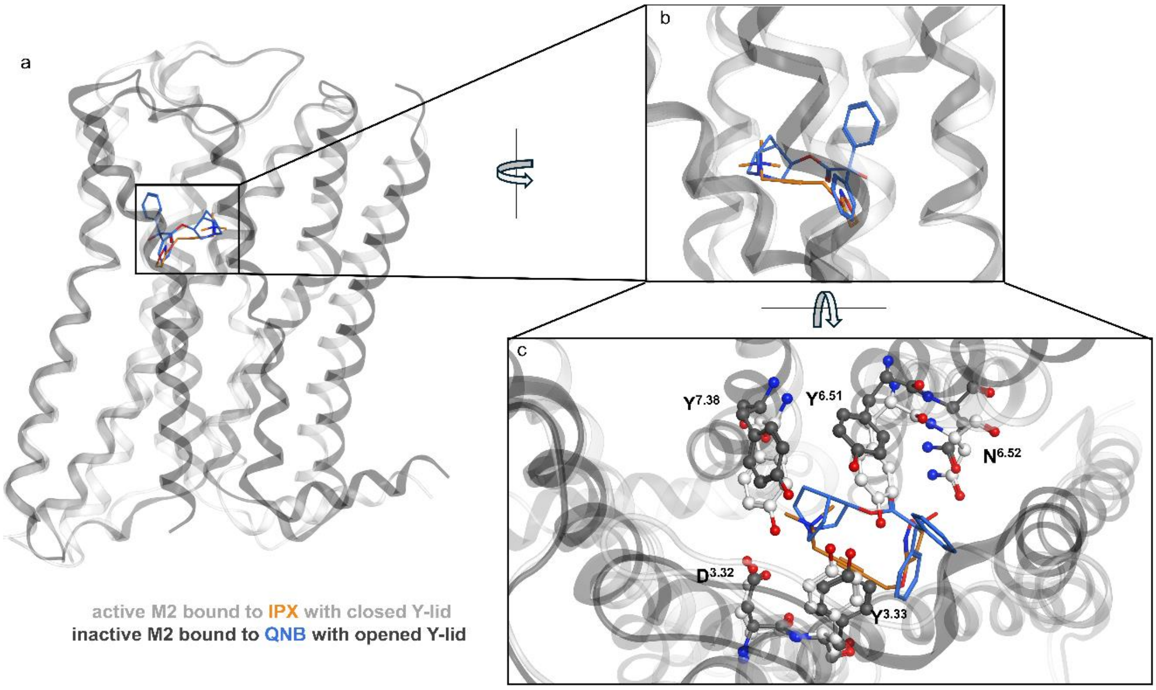


**Figure S1.** Contraction of the orthosteric binding pocket (OBP) during receptor activation. (**a**) Structural overlay of active (white) and inactive (dark grey) $M_2$ bound to the superagonist IPX (orange) and the inverse agonist QNB (blue). (**b**) Close-up on the orthosteric binding pocket. (**c**) View from above on the tyrosine lid constituted by $Y^{3.32}$, $Y^{6.51}$ and $Y^{7.38}$. During receptor activation the Y-lid closes (white residues) which contributes to the contraction of the OBP.

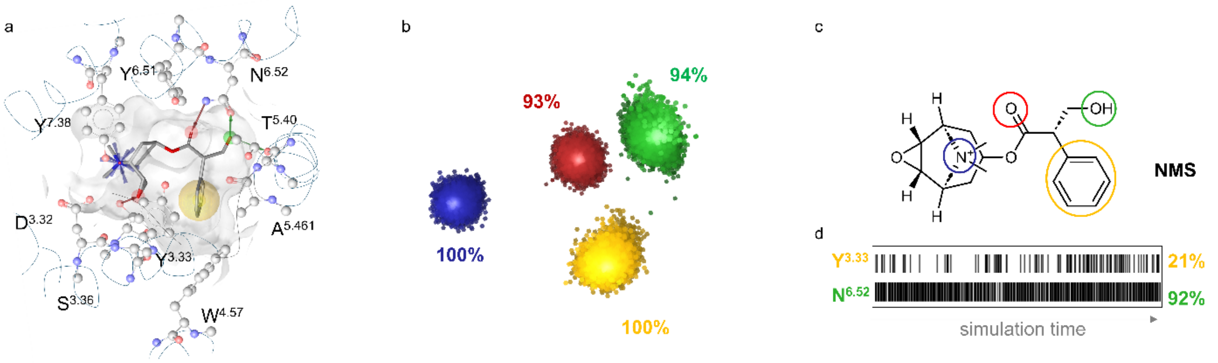


**Figure S2.** Static (**a**) and dynamic (**b**) pharmacophore of N-methylscopolamine (NMS) bound to $M_2$ (pdb-entry: 5zkc). Feature definitions: yellow – lipophilic contacts, blue – positive ionizable regions, green – H-bond-donor, red – H-bond acceptor atoms. Analogously to Figure 2 and 3. 2D structure of NMS (**c**). Functionalities corresponding to the dynamic interaction patterns in (b) were marked in the same color. (**d**) Illustration of the corresponding barcodes of lipophilic contacts between NMS's phenyl group and $Y^{3.33}$ and H-bond donor interactions, between the NMS's hydroxyl group and $N^{6.52}$, which are located on opposite sides of the OBP. Unlike all other ligands investigated in this study, NMS does not interact with $Y^{6.51}$ during the MD simulations.

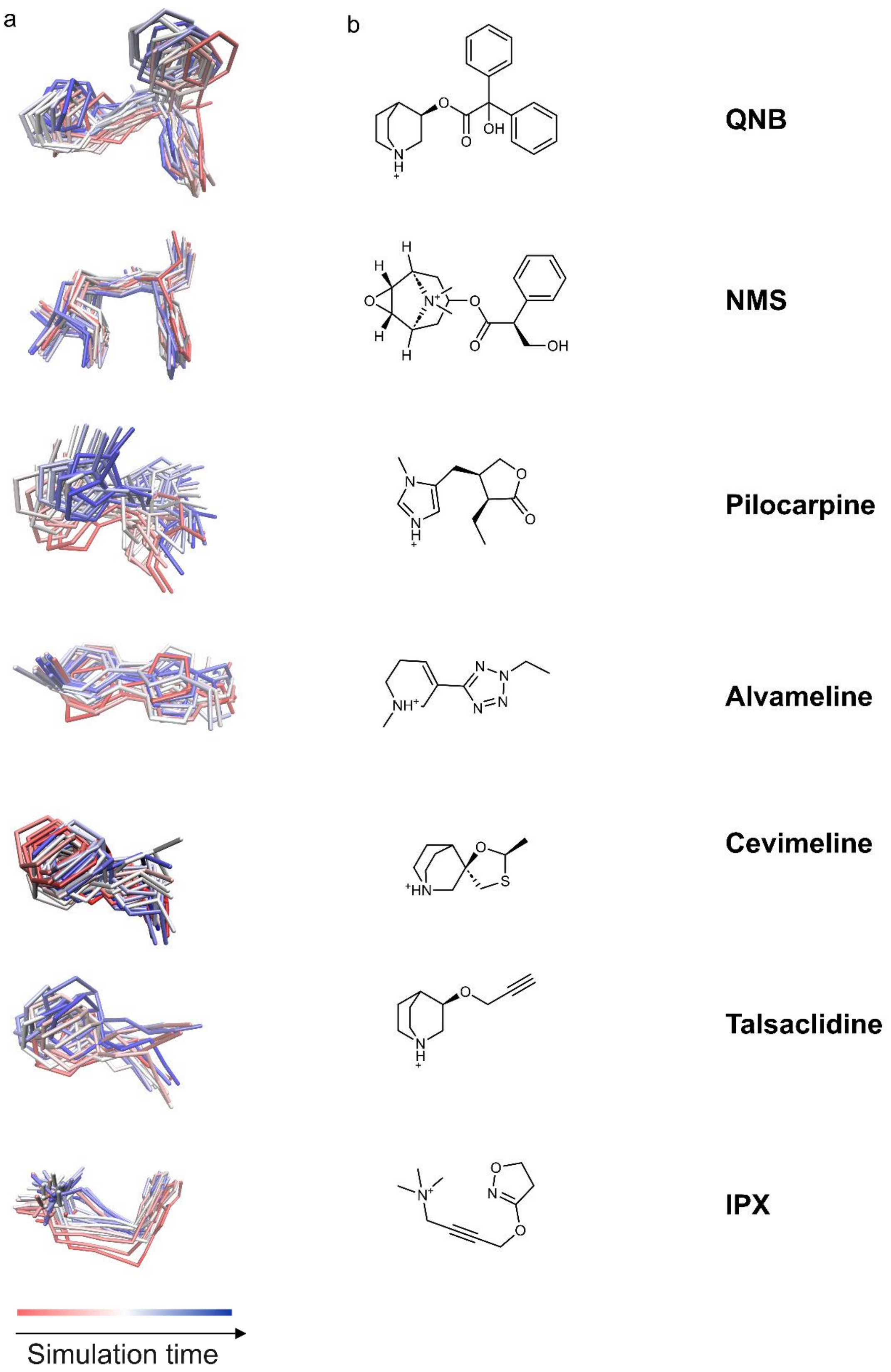


**Figure S3.** (**a**) Superposition of every 1000th frame of the different ligands throughout the trajectories. (**b**) 2D structures of orthosteric ligands used in this study. Abbreviations: QNB – 3-Quinuclidinyl benzilate; NMS – N-Methylscopolamine; IPX – Iperoxo. QNB, NMS and IPX, were crystallized in complex with $M_2$. Initial movements in the $M_2$ partial agonist complexes also result from an adaptation of OBP sidechains to the bound partial agonist. Whereby the lower number of partial agonists in X-ray and cryo-EM structures might also result from a higher heterogeneity of stabilized receptor conformations.

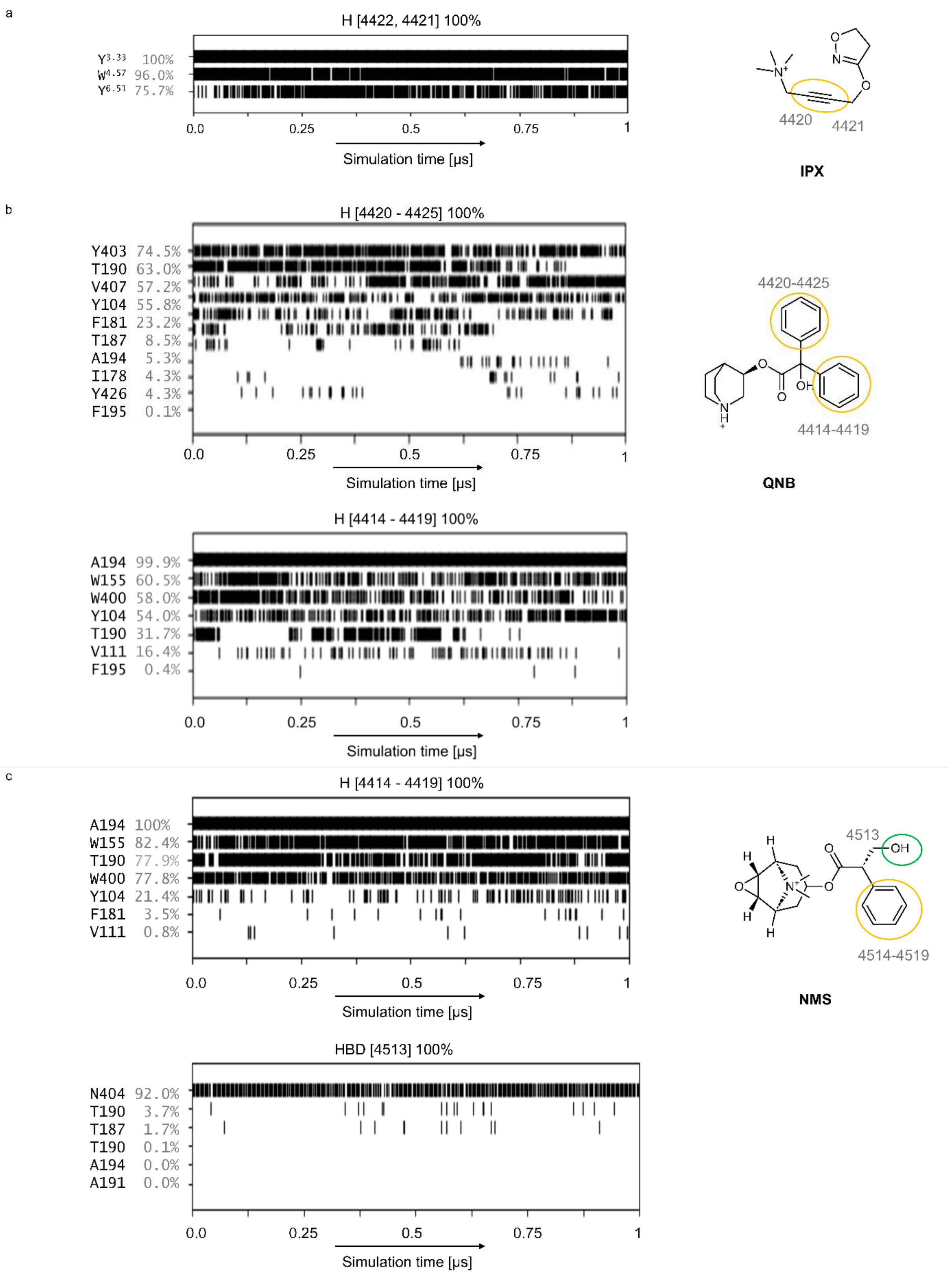


**Figure S4. Barcodes corresponding to the lipophilic features of IPX, QNB and NMS as shown in Figure 2 and S2.** (**a**) Compared to the other ligands, the lipophilic interacting group of IPX is located at a different position of the OBP. As the ligand induces a strong OBP contraction, the ethynyl moiety is able to interact with Y104$^{3.33}$ and Y403$^{6.51}$ simultaneously. (**b**) Lipophilic interactions of both phenyl moieties of QNB. (**c**) Lipophilic interactions of the phenyl moiety of NMS and H-bond donor interactions of the primary alcohol. Abbreviations: H - lipophilic interactions, HBD – H-bond donor interactions. F$^{ECL2}$ corresponds to F181. I$^{ECL2}$ corresponds to I178.

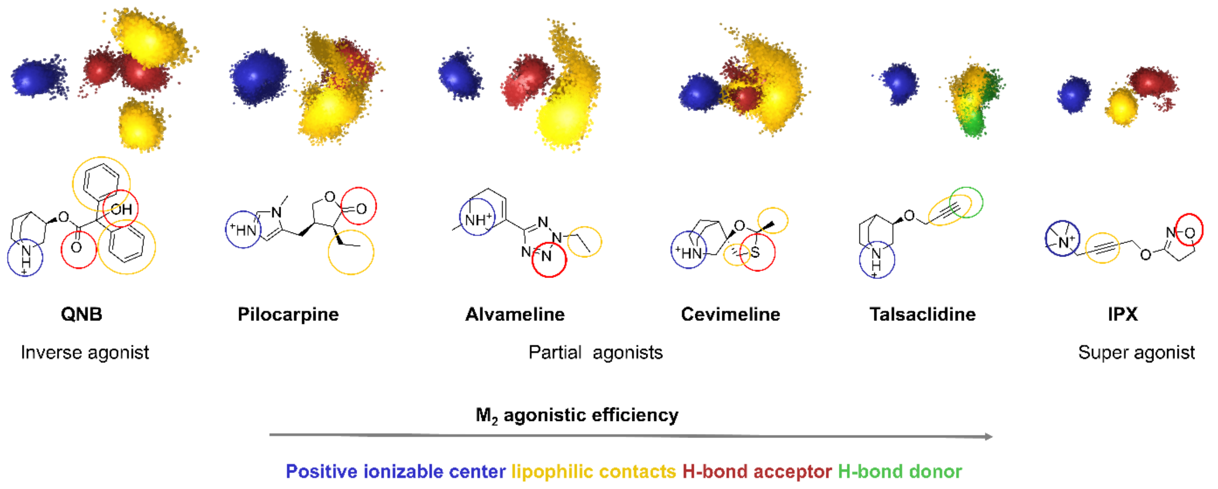


**Figure S5.** Summary of the dynamic interaction patterns correlated with the ligands' efficacy.

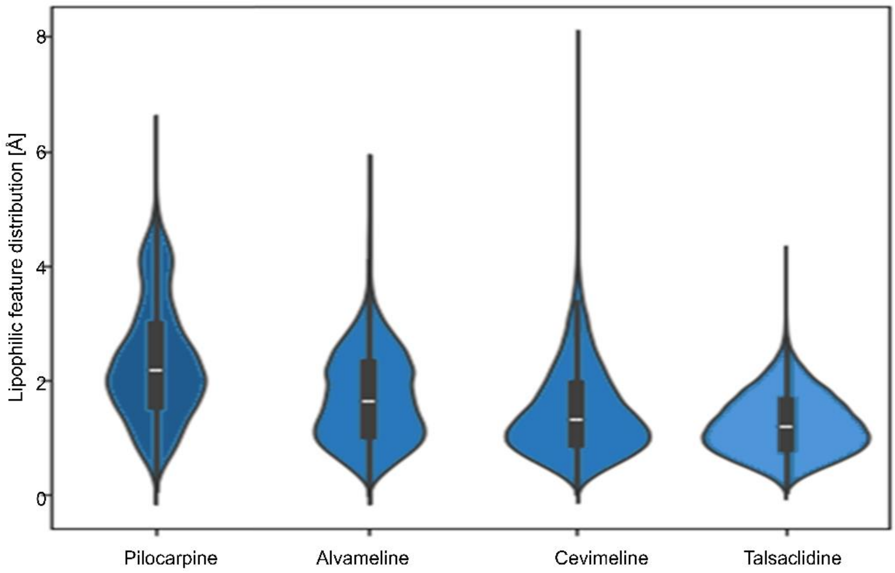


**Figure S6. Distance distribution of lipophilic interaction patterns correlates with the partial agonistic efficacy.** The distance was measured between each lipophilic feature point and the feature centroid and plotted as violin plots.

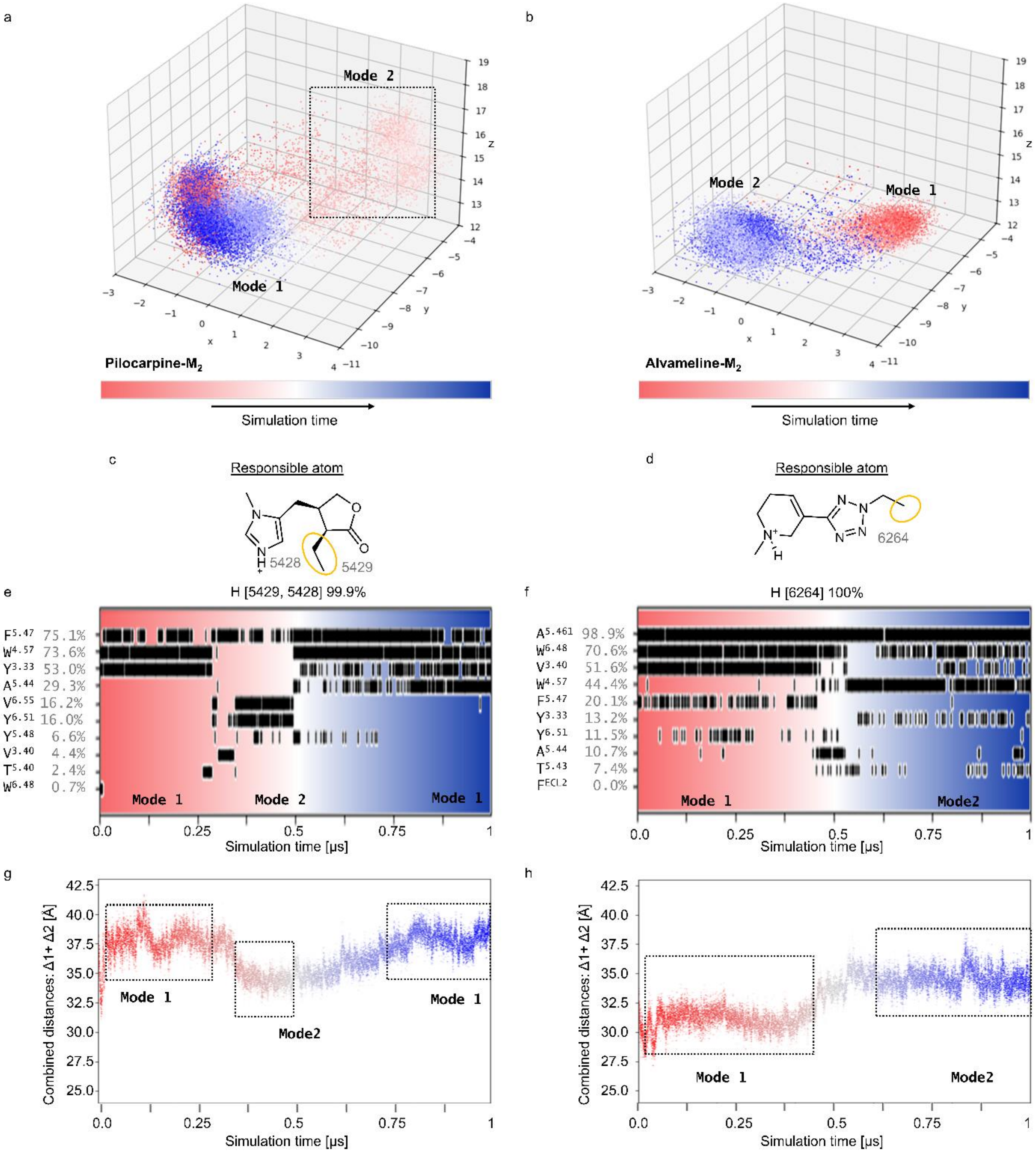


**Figure S7. Time-dependent visualization of the interaction patterns of Pilocarpine and Alvameline reveals a change in the orientation of the lipophilic moiety of the ligand that is accompanied by changes in the ligands' interaction patterns.** Both ligands adopt two main binding modes which are separated by transition states. Lipophilic interaction patterns of (**a**) Pilocarpine and (b) Alvameline colored in a time-dependent manner. The panels (c) and (d) show the corresponding barcodes. Numbers above the barcodes indicate the atom numbers marked in the corresponding ligand scaffold in (e) and (f). Abbreviations: H – van der Waals interactions. Combined distances to capture the contraction of the OBP as shown in Figure S9 are shown for M2-Pilocarpine (g) and M2-Alvameline (h). Binding mode changes are accompanied with changes in the OBP contraction, which also shows intermediate states. $F^{ECL2}$ corresponds to F181.

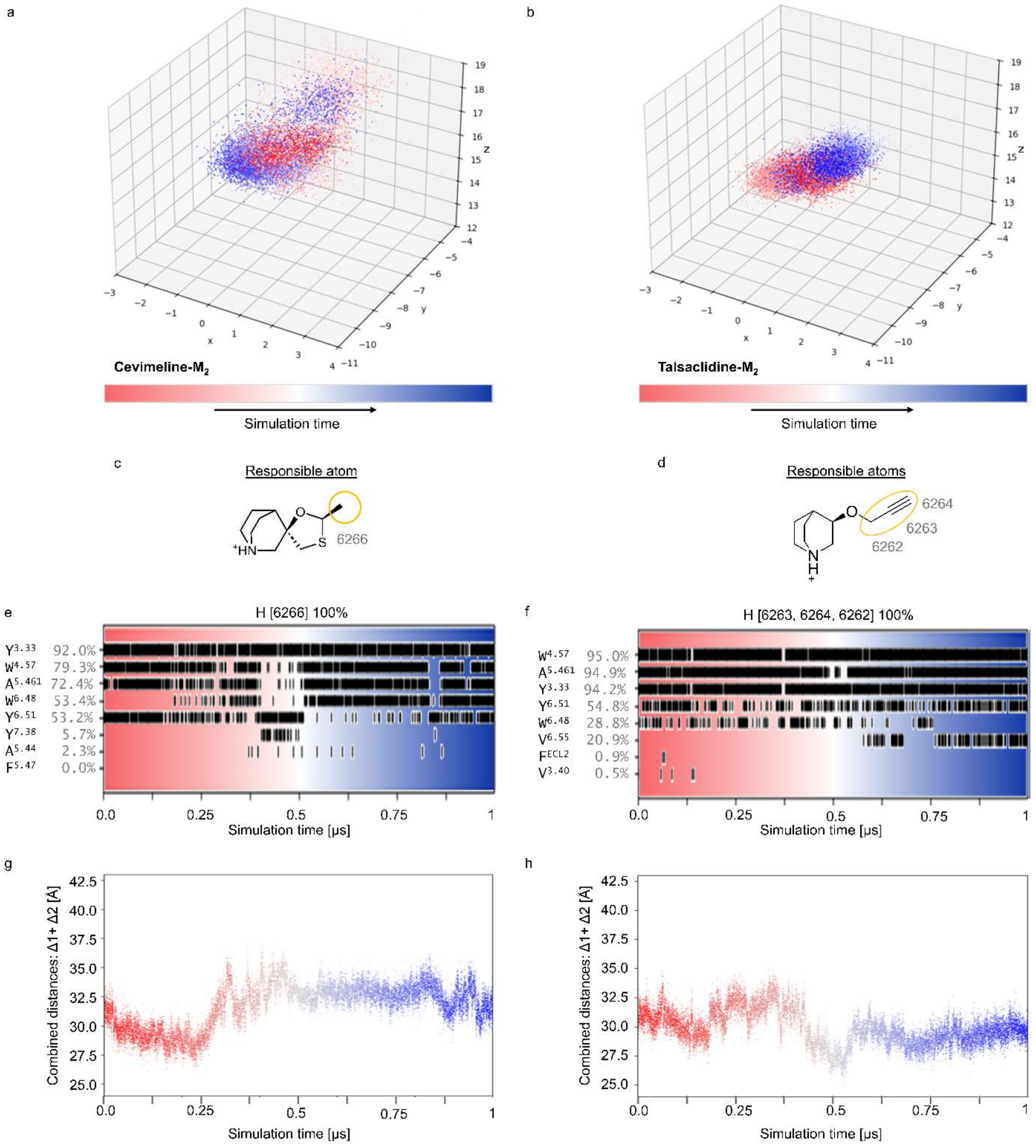


**Figure S8. Time-dependent visualization of the interaction patterns of Cevimeline and Talsaclidine.** Unlike Pilocarpine and Alvameline binding modes are optically inseparable. The spatial proximity causes more persistent interactions and only slight, fluid changes in the interacting amino acids during binding mode alterations. Lipophilic interaction patterns of (a) Cevimeline and (b) Talsaclidine colored in a time-dependent manner. The panels (c) and (d) show the corresponding barcodes. Numbers above the barcodes indicate the atom numbers marked in the corresponding ligand scaffold in (e) and (f). Abbreviations: H – van der Waals interactions. Combined distances to capture the contraction of the OBP as shown in figure 3.9 are shown for M2-Cevimeline (g) and M2-Talsaclidine (h). Interestingly, decreasing interactions with W400$^{6.48}$ seem to correlate with decreased distances through the OBP.

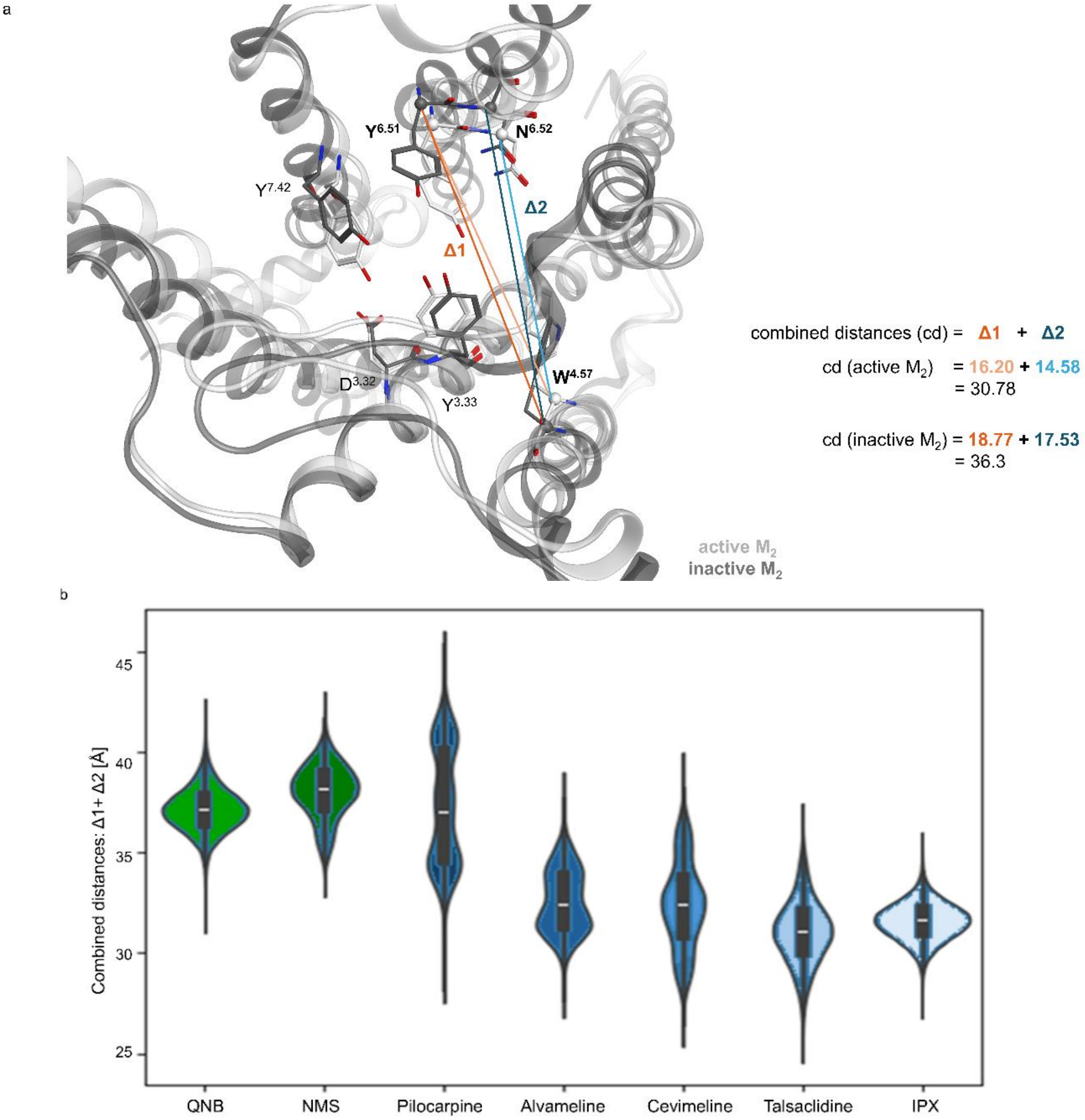


**Figure S9**. Combined distances were used as surrogate to measure $M_2$ OBP contraction. (a) Overlay of the active (white, 4MQS) and inactive (grey, 3UON) $M_2$ receptor. Distance 1 (Δ1) was measured between Cα of W155$^{4.57}$ and Cα of Y403$^{6.51}$ and is represented as dark orange line in the inactive and as light orange line in the active $M_2$ receptor. Distance 2 ((Δ2) was measured between Cα of W155$^{4.57}$ and Cα of N404$^{6.52}$ and is represented as dark blue line in the inactive and as light blue line in the active $M_2$ receptor. (b) Combined distances as represented in (a) were measured throughout the MD simulations and are represented as violin plots for each triplicate. The surrogate parameter unravels a smaller distance for the active IPX bound, than for the inactive QNB or NMS bound $M_2$ receptor. The bell-shaped distribution for QNB, NMS and IPX unravel that these ligands strongly stabilize the respective $M_2$ conformation, while the partial agonists lead to more heterogeneous OBP conformations (distance distributions).

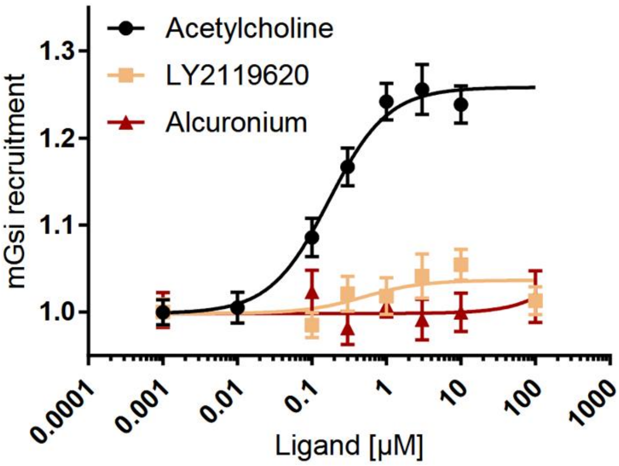


**Figure S10.** Effects of increasing concentrations of the Ago-PAM (LY2119620) and the NAM (Alcuronium) on mini Gsi recruitment by $M_2$. LY2119620 alone leads to a slight receptor activation.

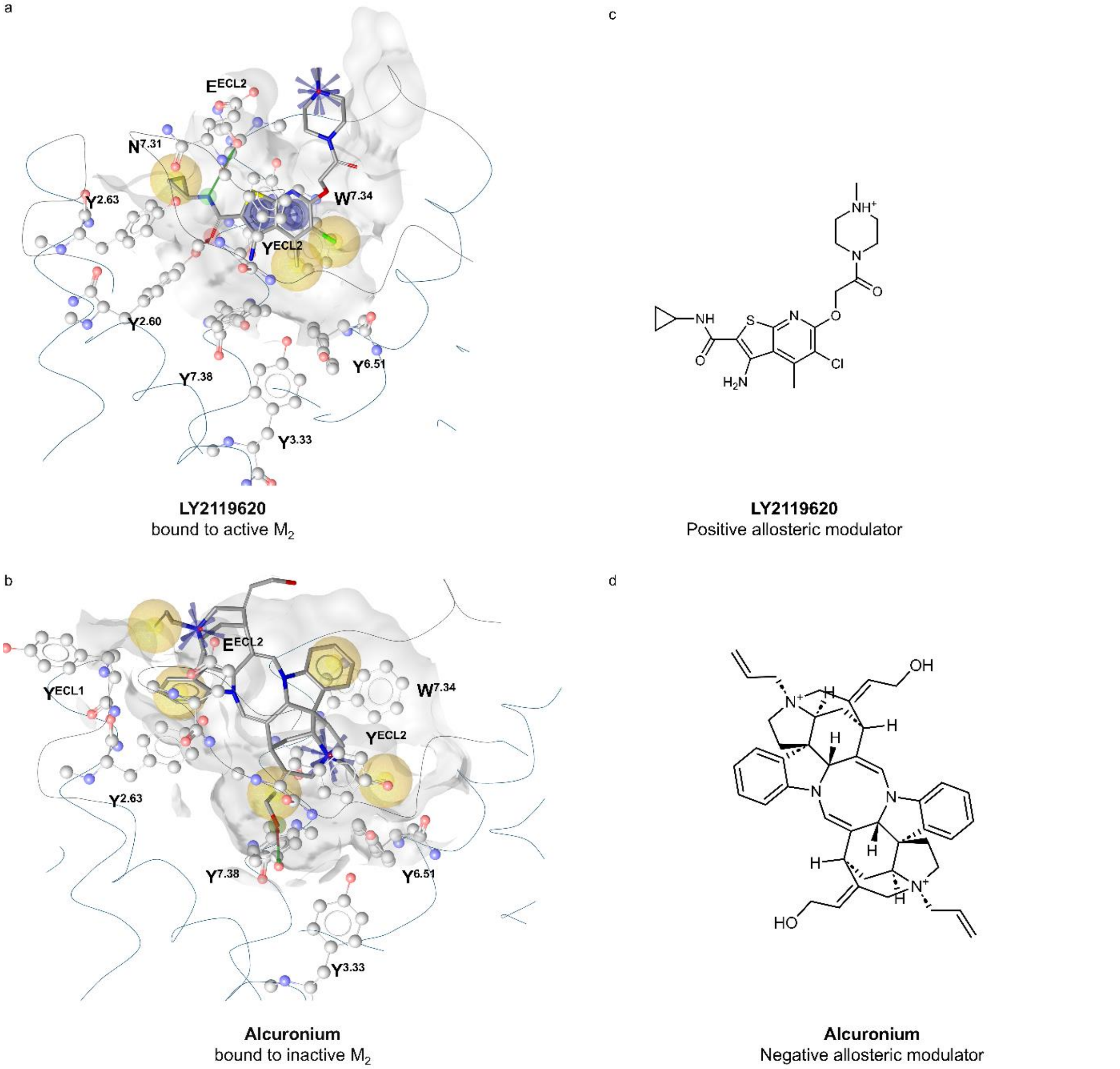


**Figure S11.** The positive (**a**) and negative (**b**) allosteric modulators LY2119620 and Alcuronium bound to the extracellular vestibule of $M_2$. The respective 2D structures are shown in (**c**) and (**d**).

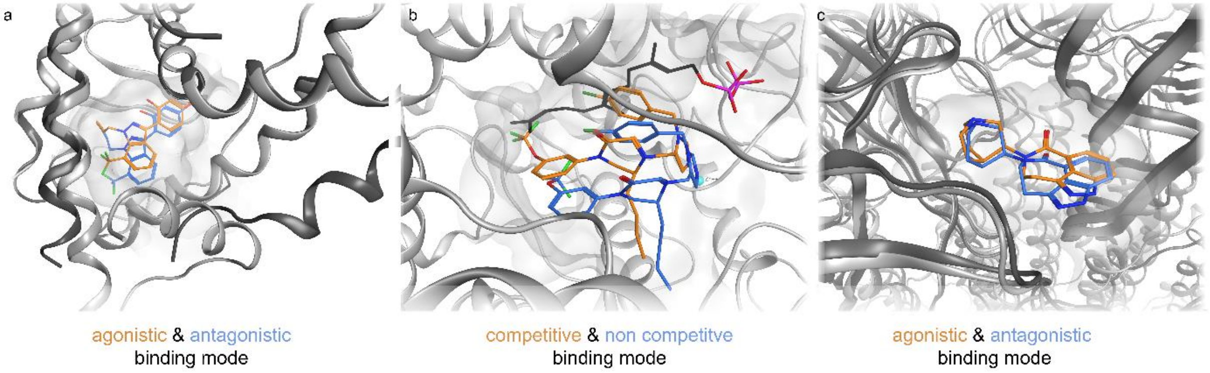


**Figure S12. Literature examples for multiple binding modes of one ligand corresponding to alternate protein states and thus, a different functional outcome.** Panel (**a**) shows the estrogen receptor α (ERα) ligand binding domain (LBD) bound to partial agonist WAY-169916. The ligand adopts four different binding modes (one in the active (WAY-169916 in orange and ERα-LBD in light grey; pdb: 2QZO) and three in the inactive ERα LBD (dark grey), we show binding mode 2 from Bruning et al. (3OS9)[21] (**b**) shows the farnesyltransferase from *Cryptococcus neoformans* bound to a mixed competitive and non-competitive inhibitor 2e. The competitive binding mode (orange) of 2e competes with the enzyme's substrate farnesyl-pyrophosphate (FPP), shown in black; the non-competitive conformation (blue) binds to the enzyme-substrate complex (pdb: 7T0C).[22] (**c**) the 5-$HT_3$ ion channel partial agonist SML is capable of stabilizing the pre activated (SML in blue, 5-$HT_3$ in dark grey, 8FRX) and the open like (SML in orange, 5-$HT_3$ in light grey, 8FSP) conformation of the ion channel.[23]

**Table S1. In vitro data of different partial agonists investigated throughout this study.** While Heinrich et al. determined potency and efficacy by measuring cAMP decrease as response to Gi coupling[24], Liu et al. used a GTPy[$S^{35}$] assay[25]. Heinz et al. determined $EC_{50}$ and $E_{max}$ in a NanoBret miniGsi recruitment assay.[19] $E_{max}$ values from Bock et al.[26] were determined by measuring dynamic mass redistribution (DMR) and normalized setting IPX as 100%. Inhibition of Ach mediated $Ca^{2+}$ mobilization by NMS was shown by Gregory et al.[27] As QNB is listed as combat agent restricting its availability, comprehensive functional data is not available.[8] Schrage et al.[28] reported binding affinities measured by replacement with radiolabeled IPX. Apart from the DMR data from Bock et al. all efficacy values were determined by setting the $E_{max}$ Acetylcholine *binding at $M_2$ as 100%.* $E_{max}$ values shown for IPX was determined by Xu et al. 2023 in a β-arrestin 2 recruitment assay.[29]

| | **$Ec_{50}$ (nM)** | **% efficacy** | **pKi-1** | **% Emax DMR Gi protein signaling** | **% $E_{max}$ DMR Gs protein signaling** |
|---|---|---|---|---|---|
| **Pilocarpine** | 1,000[19] | 39[19] | 6.54[28] | 65[26] | 8[26] |
| **Alvameline (LU25109)** | 296 +- 5.7[25] | 49.2 +- 1.6[25] | | | |
| **Cevimeline (SNI-2011)** | 1040[24] | 84[24] | | | |
| **Talsaclidine (WAL-2014)** | 1535[24];<br>849 +- 69[25] | 92[24];<br>94.4 +- 2.8[25] | | | |
| **NMS** | | | 9.26[28] | | |
| **IPX** | | 151[29] | 10.23[28] | 100[26] | 100[26] |

**Table S2. M$_2$-RMSDs**. Alignment to the backbone heavy atoms of frame 1; dynamics replicate: 1, 2, 3

M$_2$-Iperoxo

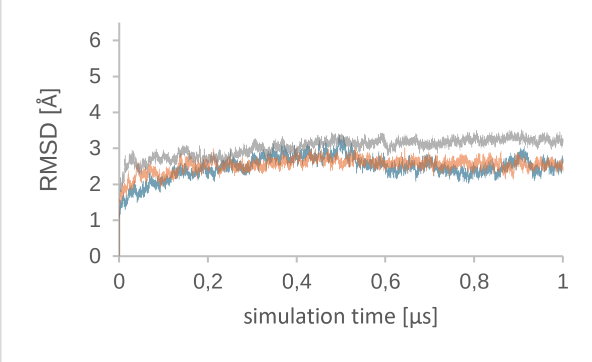


M$_2$-Alvameline

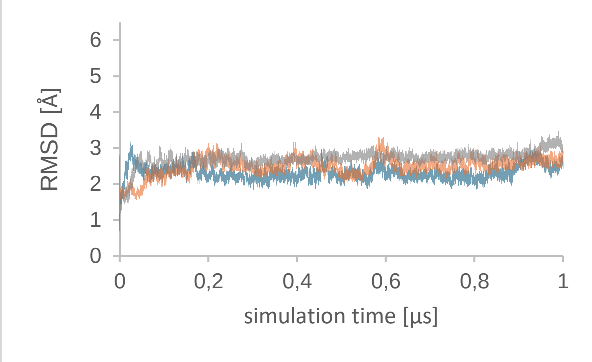


M$_2$-Cevimeline

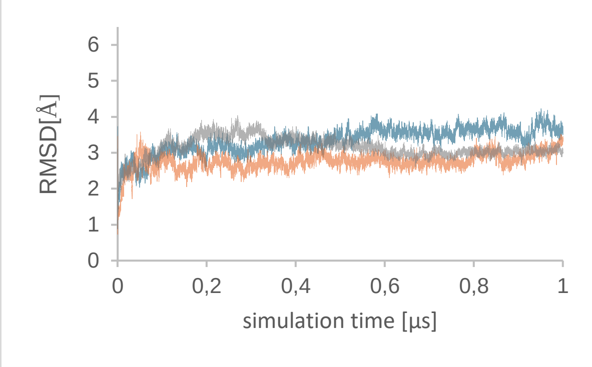


M$_2$-Talsaclidine

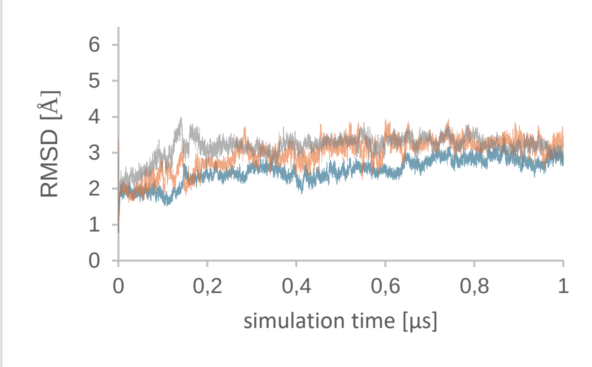


M$_2$-Pilocarpine

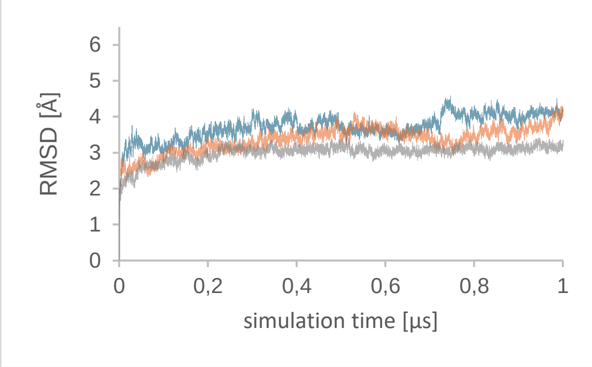


M$_2$-Quinuclidinylbenzilate

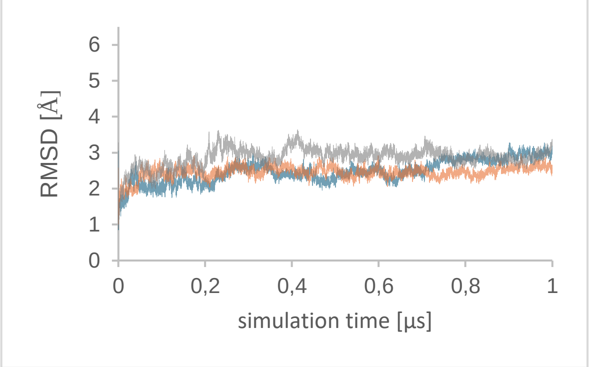


M$_2$-N-Methylscopolamine

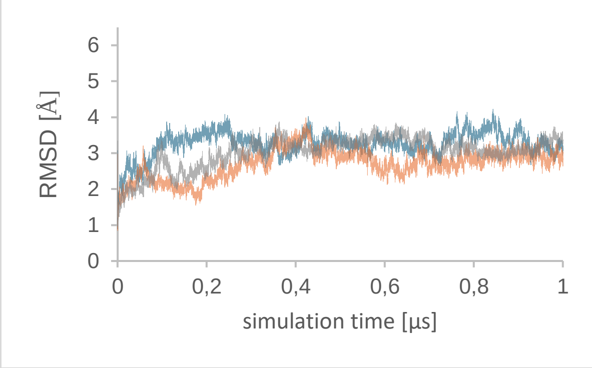

**Table S3. Ligand RMSDs.** Alignment to the ligand heavy atoms of frame 1; dynamics replicate: 1, 2, 3

M2-Iperoxo

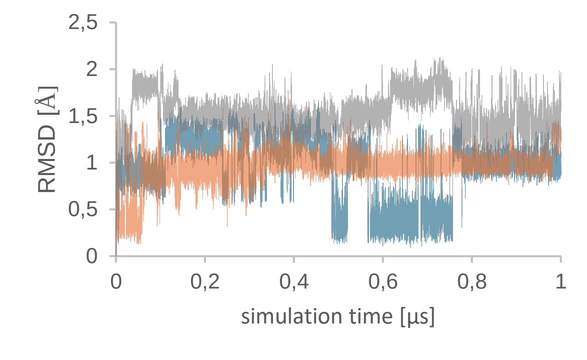


M2-Alvameline NR

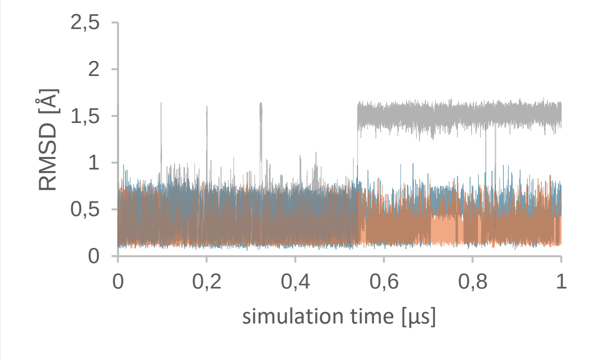


M2-Cevimeline

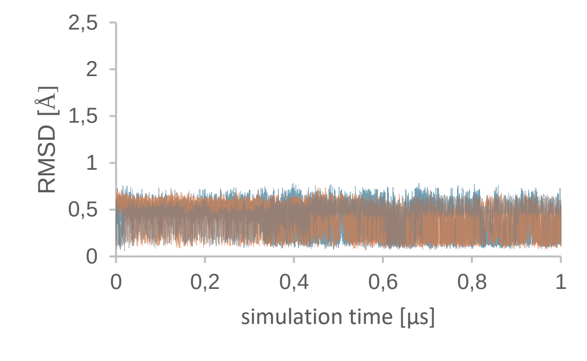


M2-Talsaclidine

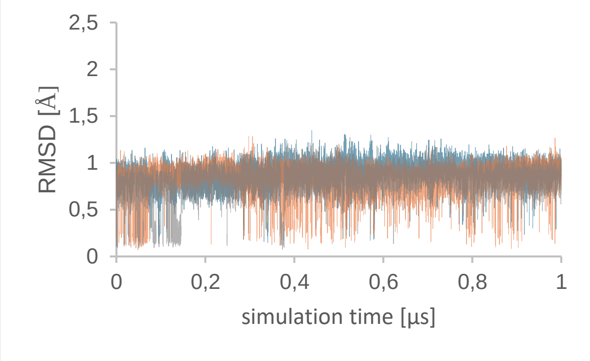


M2-Pilocarpine

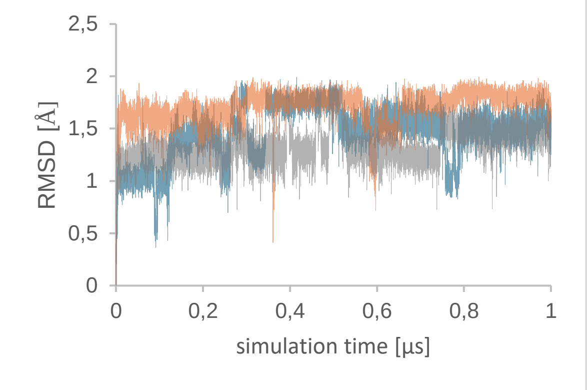


M2-Quinuclidinylbenzilate

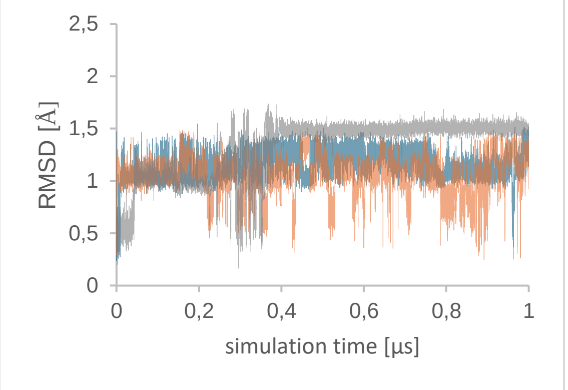


M2-N-Methylscopolamine

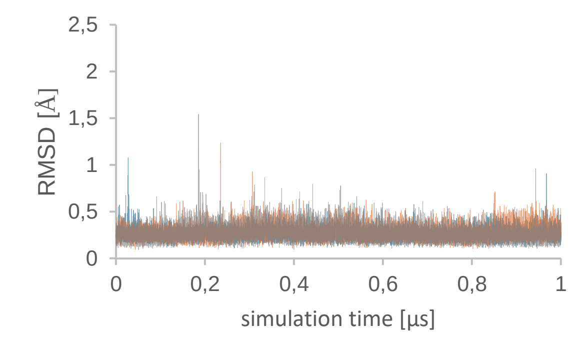

### References


(1) Chemical Computing Group ULC, 910-1010 Sherbrooke St. W., Montreal, QC H3A 2R7. *Molecular Operating Environment (MOE), 2024.0601*, 2020.

(2) Kruse, A. C.; Ring, A. M.; Manglik, A.; Hu, J.; Hu, K.; Eitel, K.; Hübner, H.; Pardon, E.; Valant, C.; Sexton, P. M.; Christopoulos, A.; Felder, C. C.; Gmeiner, P.; Steyaert, J.; Weis, W. I.; Garcia, K. C.; Wess, J.; Kobilka, B. K. Activation and allosteric modulation of a muscarinic acetylcholine receptor. *Nature* **2013**, *504* (7478), 101–106. DOI: 10.1038/nature12735. Published Online: Nov. 20, 2013.

(3) Jones, G.; Willett, P.; Glen, R. C.; Leach, A. R.; Taylor, R. Development and validation of a genetic algorithm for flexible docking. *Journal of molecular biology* **1997**, *267* (3), 727–748. DOI: 10.1006/jmbi.1996.0897.

(4) Evers, A.; Hessler, G.; Matter, H.; Klabunde, T. Virtual screening of biogenic amine-binding G-protein coupled receptors: comparative evaluation of protein- and ligand-based virtual screening protocols. *Journal of Medicinal Chemistry* **2005**, *48* (17), 5448–5465. DOI: 10.1021/jm050090o.

(5) Verdonk, M. L.; Cole, J. C.; Hartshorn, M. J.; Murray, C. W.; Taylor, R. D. Improved protein-ligand docking using GOLD. *Proteins* **2003**, *52* (4), 609–623. DOI: 10.1002/prot.10465.

(6) Wolber, G.; Langer, T. LigandScout: 3-D Pharmacophores Derived from Protein-Bound Ligands and Their Use as Virtual Screening Filters. *Journal of Chemical Information and Modeling* **2005**, *45* (1), 160–169. DOI: 10.1021/ci049885e.

(7) Wolber, G.; Dornhofer, A. A.; Langer, T. Efficient overlay of small organic molecules using 3D pharmacophores. *Journal of Computer-Aided Molecular Design* **2006**, *20* (12), 773–788. DOI: 10.1007/s10822-006-9078-7.

(8) Haga, K.; Kruse, A. C.; Asada, H.; Yurugi-Kobayashi, T.; Shiroishi, M.; Zhang, C.; Weis, W. I.; Okada, T.; Kobilka, B. K.; Haga, T.; Kobayashi, T. Structure of the human M2 muscarinic acetylcholine receptor bound to an antagonist. *Nature* **2012**, *482* (7386), 547–551. DOI: 10.1038/nature10753. Published Online: Jan. 25, 2012.

(9) Suno, R.; Lee, S.; Maeda, S.; Yasuda, S.; Yamashita, K.; Hirata, K.; Horita, S.; Tawaramoto, M. S.; Tsujimoto, H.; Murata, T.; Kinoshita, M.; Yamamoto, M.; Kobilka, B. K.; Vaidehi, N.; Iwata, S.; Kobayashi, T. Structural insights into the subtype-selective antagonist binding to the M2 muscarinic receptor. *Nature chemical biology* **2018**, *14* (12), 1150–1158. DOI: 10.1038/s41589-018-0152-y. Published Online: Nov. 12, 2018.

(10) *Schrödinger Release 2021-1*:*Maestro, Schrödinger*, 2021.

(11) Zielkiewicz, J. Structural properties of water: comparison of the SPC, SPCE, TIP4P, and TIP5P models of water. *The Journal of chemical physics* **2005**, *123* (10), 104501. DOI: 10.1063/1.2018637.

(12) Lomize, M. A.; Pogozheva, I. D.; Joo, H.; Mosberg, H. I.; Lomize, A. L. OPM database and PPM web server: resources for positioning of proteins in membranes. *Nucleic Acids Res* **2012**, *40* (D1), D370-D376. DOI: 10.1093/nar/gkr703.

(13) Jorgensen, W. L.; Maxwell, D. S.; Tirado-Rives, J. Development and Testing of the OPLS All-Atom Force Field on Conformational Energetics and Properties of Organic Liquids. *Journal of the American Chemical Society* **1996**, *118* (45), 11225–11236. DOI: 10.1021/ja9621760.

(14) Bowers, K. J.; Chow, D. E.; Xu, H.; Dror, R. O.; Eastwood, M. P.; Gregersen, B. A.; Klepeis, J. L.; Kolossvary, I.; Moraes, M. A.; Sacerdoti, F. D.; Salmon, J. K.; Shan,

Y.; Shaw, D. E. Scalable Algorithms for Molecular Dynamics Simulations on Commodity Clusters. In *ACM/IEEE SC 2006 Conference (SC'06)*; IEEE, 2006 - 2006; p 43. DOI: 10.1109/SC.2006.54.
(15) Humphrey, W.; Dalke, A.; Schulten, K. VMD: Visual molecular dynamics. *Journal of Molecular Graphics* **1996**, *14* (1), 33–38. DOI: 10.1016/0263-7855(96)00018-5.
(16) Schaller, D.; Šribar, D.; Noonan, T.; Deng, L.; Nguyen, T. N.; Pach, S.; Machalz, D.; Bermudez, M.; Wolber, G. Next generation 3D pharmacophore modeling. *WIREs Comput Mol Sci* **2020**, *10* (4), e1468. DOI: 10.1002/wcms.1468.
(17) Wu, B.; Chien, E. Y. T.; Mol, C. D.; Fenalti, G.; Liu, W.; Katritch, V.; Abagyan, R.; Brooun, A.; Wells, P.; Bi, F. C.; Hamel, D. J.; Kuhn, P.; Handel, T. M.; Cherezov, V.; Stevens, R. C. Structures of the CXCR4 chemokine GPCR with small-molecule and cyclic peptide antagonists. *Science (New York, N.Y.)* **2010**, *330* (6007), 1066–1071. DOI: 10.1126/science.1194396. Published Online: Oct. 7, 2010.
(18) Pedregosa, F.; Varoquaux, G.; Gramfort, A.; Michel, V.; Thirion, B.; Grisel, O.; Blondel, M.; Prettenhofer, P.; Weiss, R.; Dubourg, V.; Vanderplas, V.; Passos, A.; Cournapeau, D.; Brucher, M.; Perrot, M.; Duchesnay, É. Scikit-learn: Machine learning in Python. *Journal of Machine Learning Research* **2011** (12), 2825–2830.
(19) Heinz, C. S.; Bermudez, M.; Jaiswal, N.; Große, C.; Kauk, M.; Hoffmann, C.; Holzgrabe, U. Hybridization into a Bitopic Ligand Increased Muscarinic Receptor Activation for Isopilocarpine but Not for Pilocarpine Derivatives. *Journal of Natural Products* **2023**, *86* (4), 869–881. DOI: 10.1021/acs.jnatprod.2c01079.
(20) Leonhardt, J.; Haider, R. S.; Sponholz, C.; Leonhardt, S.; Drube, J.; Spengler, K.; Mihaylov, D.; Neugebauer, S.; Kiehntopf, M.; Lambert, N. A.; Kortgen, A.; Bruns, T.; Tacke, F.; Hoffmann, C.; Bauer, M.; Heller, R. Circulating Bile Acids in Liver Failure Activate TGR5 and Induce Monocyte Dysfunction. *Cellular and Molecular Gastroenterology and Hepatology* **2021**, *12* (1), 25–40. DOI: 10.1016/j.jcmgh.2021.01.011.
(21) Bruning, J. B.; Parent, A. A.; Gil, G.; Zhao, M.; Nowak, J.; Pace, M. C.; Smith, C. L.; Afonine, P. V.; Adams, P. D.; Katzenellenbogen, J. A.; Nettles, K. W. Coupling of receptor conformation and ligand orientation determine graded activity. *Nature chemical biology* **2010**, *6* (11), 837–843. DOI: 10.1038/nchembio.451.
(22) Wang, Y.; Xu, F.; Nichols, C. B.; Shi, Y.; Hellinga, H. W.; Alspaugh, J. A.; Distefano, M. D.; Beese, L. S. Structure-Guided Discovery of Potent Antifungals that Prevent Ras Signaling by Inhibiting Protein Farnesyltransferase. *Journal of Medicinal Chemistry* **2022**, *65* (20), 13753–13770. DOI: 10.1021/acs.jmedchem.2c00902.
(23) Felt, K.; Stauffer, M.; Salas-Estrada, L.; Guzzo, P. R.; Xie, D.; Huang, J.; Filizola, M.; Chakrapani, S. Structural basis for partial agonism in 5-HT3A receptors. *Nature Structural & Molecular Biology* **2024**, *31* (4), 598–609. DOI: 10.1038/s41594-023-01140-2.
(24) Heinrich, J. N.; Butera, J. A.; Carrick, T.; Kramer, A.; Kowal, D.; Lock, T.; Marquis, K. L.; Pausch, M. H.; Popiolek, M.; Sun, S.-C.; Tseng, E.; Uveges, A. J.; Mayer, S. C. Pharmacological comparison of muscarinic ligands: historical versus more recent muscarinic M1-preferring receptor agonists. *European journal of pharmacology* **2009**, *605* (1-3), 53–56. DOI: 10.1016/j.ejphar.2008.12.044.
(25) Liu, B.; Croy, C. H.; Hitchcock, S. A.; Allen, J. R.; Rao, Z.; Evans, D.; Bures, M. G.; McKinzie, D. L.; Watt, M. L.; Stuart Gregory, G.; Hansen, M. M.; Hoogestraat, P. J.; Jamison, J. A.; Okha-Mokube, F. M.; Stratford, R. E.; Turner, W.; Bymaster, F.; Felder, C. C. Design and synthesis of N-[6-(Substituted Aminoethylideneamino)-2-

Hydroxyindan-1-yl]arylamides as selective and potent muscarinic M1 agonists. *Bioorganic & medicinal chemistry letters* **2015**, *25* (19), 4158–4163. DOI: 10.1016/j.bmcl.2015.08.011.
(26) Bock, A.; Merten, N.; Schrage, R.; Dallanoce, C.; Bätz, J.; Klöckner, J.; Schmitz, J.; Matera, C.; Simon, K.; Kebig, A.; Peters, L.; Müller, A.; Schrobang-Ley, J.; Tränkle, C.; Hoffmann, C.; Amici, M. de; Holzgrabe, U.; Kostenis, E.; Mohr, K. The allosteric vestibule of a seven transmembrane helical receptor controls G-protein coupling. *Nature communications* **2012**, *3*, 1044. DOI: 10.1038/ncomms2028.
(27) Gregory, K. J.; Hall, N. E.; Tobin, A. B.; Sexton, P. M.; Christopoulos, A. Identification of Orthosteric and Allosteric Site Mutations in $M_2$ Muscarinic Acetylcholine Receptors That Contribute to Ligand-selective Signaling Bias * . *Journal of Biological Chemistry* **2010**, *285* (10), 7459–7474. DOI: 10.1074/jbc.M109.094011.
(28) Schrage, R.; Holze, J.; Klöckner, J.; Balkow, A.; Klause, A. S.; Schmitz, A.-L.; Amici, M. de; Kostenis, E.; Tränkle, C.; Holzgrabe, U.; Mohr, K. New insight into active muscarinic receptors with the novel radioagonist [3H]iperoxo. *Biochemical Pharmacology* **2014**, *90* (3), 307–319. DOI: 10.1016/j.bcp.2014.05.012.
(29) Xu, J.; Wang, Q.; Hübner, H.; Hu, Y.; Niu, X.; Wang, H.; Maeda, S.; Inoue, A.; Tao, Y.; Gmeiner, P.; Du, Y.; Jin, C.; Kobilka, B. K. Structural and dynamic insights into supra-physiological activation and allosteric modulation of a muscarinic acetylcholine receptor. *Nature communications* **2023**, *14* (1), 376. DOI: 10.1038/s41467-022-35726-z. Published Online: Jan. 23, 2023.
